# Comparable genomic copy number aberrations differ across astrocytoma malignancy grades

**DOI:** 10.1101/446831

**Authors:** Nives Pećina-Šlaus, Anja Kafka, Kristina Gotovac, Monika Logara, Anja Bukovac, Robert Bakarić, Fran Borovečki

**Author notes:** Corresponding author Nives PeCina-Slaus Laboratory of Neurooncology, Croatian Institute for Brain Research, School of Medicine University of Zagreb, Salata 3, HR-10000 Zagreb, Croatia; ORCHID iD 0000-0002-3334-7671, AnjaKafka; ORCHID iD 0000-0001-8457-5757, KristinaGotovac; ORCHID iD, Monika Logara, ORCHID iD 0000-0003-0613-693X, Anja Bukovac, ORCHID iD 0000-0001-6587-8314, Robert Bakaric, ORCHID iD 0000-0002-2102-586X, FranBorovecki, ORCHID iD 0000-0002-5178-7929.

## Abstract

Malignancy grades of astrocytomas were analyzed for copy number aberrations by aCGH. Altogether 1438 CNA were found of which losses prevailed. We searched for regions that are more likely to drive cancer pathogenesis with Bioconductor package and Genomic Identification of Significant Targets in Cancer. On our total sample significant deletions affected 14 chromosomal regions, out of which deletions at 17p13.2, 9p21.3, 13q12.11, 22q12.3 remained significant even at 0.05 q-value. When divided to malignancy groups, the regions identified as significantly deleted in high grades were: 9p21.3; 17p13.2; 10q24.2; 14q21.3; 1p36.11 and 13q12.11, while amplified were: 3q28; 12q13.3 and 21q22.3. Low grades comprised significant deletions at 3p14.3; 11p15.4; 15q15.1; 16q22.1; 20q11.22 and 22q12.3 indicating their involvement in early stages of tumorigenesis. Significantly enriched pathways brought by DAVID software were PI3K-Akt, Cytokine-cytokine receptor, NOD-like receptor, Jak-STAT, RIG-II-like receptor and Toll-like receptor pathways. HPV and herpex simplex infection and inflammation pathways were also represented. Present study brings new data to astrocytoma research amplifying the wide spectrum of changes which could help us identify the regions critical for tumorigenesis.

## Introduction

Despite the advances in astrocytoma genetics and molecular characterization, many questions about the biology of these most common primary central nervous system tumors remain unanswered. The heterogeneity of their histological features is accompanied with marked genetic and genomic heterogeneity (Appin&Brat, 2014; Abou-El-Ardat et al, 2017). However, distinct genomic patterns are emerging indicating the involvement of prominent signaling pathways, namely RTK/RAS/PI-3K, p53 and RB signaling (Cancer Genome Atlas (TCGA) Research Network, 2008; Verhaak et al, 2010; Chin et al, 2011; Pećina-Šlaus et al, 2014a; Kafka et al, 2017). Based on new discoveries on patterns of somatic mutations and DNA copy number variations involved in glioblastoma etiology four molecular signatures were proposed that classify glioblastoma into proneural, neural, classical and mesenchymal (Verhaak et al, 2010; Brennan et al, 2013).

World Health Organization (WHO) classifies astrocytomas into four grades (Louis et al, 2016; Nasser & Mehdipour, 2017) which denote their malignancy levels. Grades differ in tumor histology, growth potential, and tendency for progression, age distribution, behavior and prognosis. Astrocytomas grade I (pilocytic) typically shows benign clinical behavior and malignant progression is extraordinarily rare, describing it as a benign tumor. Astrocytomas grades II and III (diffuse and anaplastic) can progress and evolve to grade IV tumors (glioblastoma). Primary glioblastomas arise *de novo* without cognition of precursory lesions of lower grades. The highly invasive nature of glioblastoma makes it a deadly malignant tumor that is still untreatable (Paw et al, 2015).

It has been shown by several investigations that genomic copy number changes play important roles in glioblastoma (Barbashina et al, 2005; Roerig et al, 2005; Ruano et al, 2010; Beroukhim et al, 2007; Yang et al, 2011; Crespo et al, 2012). The objective of our present study was to discover genetic regions at high resolution that are altered in astrocytomas of different malignancy grades in order to identify candidate regions and genes that are appearing constantly across malignancy grades but also those that are specific for the progression of glial tumors.

Molecular mechanisms and genes involved in the formation and progression of astrocytomas is far from being understood. One of the unanswered questions is the progression of secondary glioblastoma from tumors of lower grades. Present study in which the grades of astrocytoma were compared with their genetic alterations aims to elucidate the events responsible for glioblastoma to acquire its marked malignancy. Thus we performed a genome-wide survey of gene copy number changes using array comparative genomic hybridization (aCGH), the method that has been widely used in genetic profiling of different types of cancer (Carter, 2007; Mohapatra and Sharma, 2013).

aCGH is a reliable and sensitive technique for detecting gene copy number aberration (CNA) across the entire genome. Oligonucleotide microarrays provide high resolution and diagnostic yield of detection of copy number changes comprised in the tumor genome (Banerjee, 2013; Riegel, 2014). We were interested to characterize unbalanced genomic changes gains/duplications and losses/deletions in our astrocytoma patients and offer potential candidate genes characteristic for malignancy grade as well as those recurring in several or all grades. Hence, we investigated the genomes of 14 intracranial astrocytic brain tumors of different WHO grades for changes in DNA copy number using high-resolution CGH arrays that contained 180 000 probes with the possibility to screen the genome with an average resolution of 10–50 kb. We further aimed to systematically analyze the information on CNA by bioinformatics approach utilizing rCGH (Bioconductor package) and GISTIC (Genomic Identification of Significant Targets in Cancer) 2.0.23. in order to comprehend which findings are relevant for astrocytoma biology.

## Results

### CNA in our total astrocytoma sample

The aCGH profiles demonstrated many differences in astrocytoma DNA when compared to normal control on the array. The multitude of changes that we observed in astrocytoma cells is indicative of the accumulation of deletions and amplifications characteristic of tumor cells. Astrocytoma patients showed gains and losses on many chromosomal regions. There were also a substantial number of amplifications and deletions but lower than the frequencies of the first two types of aberrations. Altogether, our aCGH results showed 1438 CNA found across astrocytomas of different malignancy grades, including 21 amplifications, 397 gains, 929 losses and 91 deletions. Losses dominated over gains and deletions over amplifications.

When grouping tumors grades I, II and III as one category and glioblastomas (grade IV) as another, we noticed that the first group predominantly harbored losses and deletions, while glioblastomas were characterized with more gains and amplifications. Average number of deletions and losses per I, II and III grouped tumors was 145 and per glioblastomas 32.8. Average number of gains and amplifications was 18.2 per first group tumors, and 36.3 per glioblastomas. Subsequent bioinformatics analyses denoted this as a visual trend that happened at higher thresholds but was not statistically relevant.

### CNA in pilocytic astrocytomas (grade I)

We assigned changes to the specific astrocytoma grade and found that patients with pilocytic astrocytomas (grade I) shared many jointly affected regions. Recurrent losses and recurrent gains are presented in Table 1. All recurrent CNA are demonstrated in more details in Table. We noticed that the number of losses (21 losses) recurring in pilocytic cases exceeded the number of recurrent gains (2 gains).

**Table 1.**
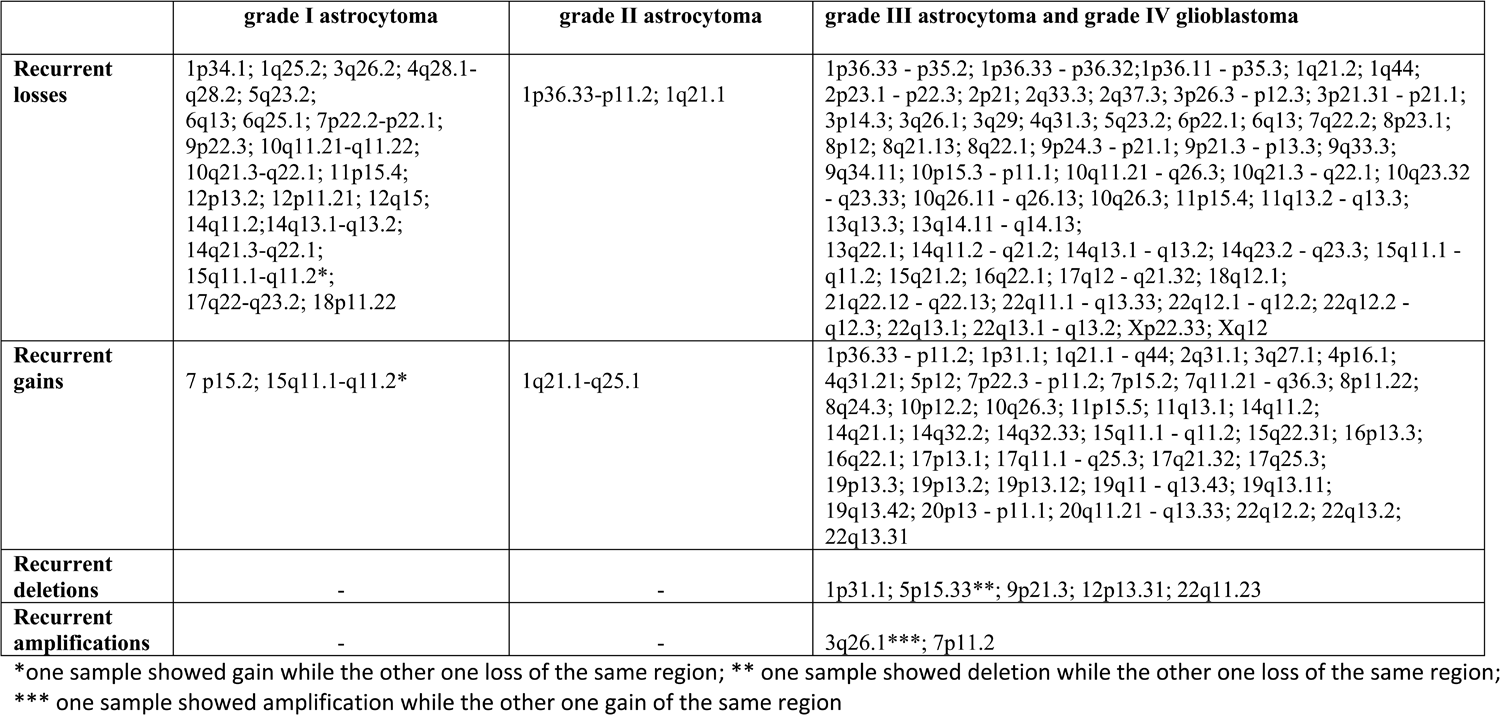
Aberrant regions shared between astrocytomas of the same malignancy grade.

|  | grade I astrocytoma | grade II astrocytoma | grade III astrocytoma and grade IV glioblastoma |
| --- | --- | --- | --- |
| **Recurrent losses** | 1p34.1; 1q25.2; 3q26.2; 4q28.1-q28.2; 5q23.2;<br>6q13; 6q25.1; 7p22.2-p22.1;<br>9p22.3; 10q11.21-q11.22;<br>10q21.3-q22.1; 11p15.4;<br>12p13.2; 12p11.21; 12q15;<br>14q11.2; 14q13.1-q13.2;<br>14q21.3-q22.1;<br>15q11.1-q11.2*;<br>17q22-q23.2; 18p11.22 | 1p36.33-p11.2; 1q21.1 | 1p36.33 - p35.2; 1p36.33 - p36.32; 1p36.11 - p35.3; 1q21.2; 1q44;<br>2p23.1 - p22.3; 2p21; 2q33.3; 2q37.3; 3p26.3 - p12.3; 3p21.31 - p21.1;<br>3p14.3; 3q26.1; 3q29; 4q31.3; 5q23.2; 6p22.1; 6q13; 7q22.2; 8p23.1;<br>8p12; 8q21.13; 8q22.1; 9p24.3 - p21.1; 9p21.3 - p13.3; 9q33.3;<br>9q34.11; 10p15.3 - p11.1; 10q11.21 - q26.3; 10q21.3 - q22.1; 10q23.32 - q23.33; 10q26.11 - q26.13; 10q26.3; 11p15.4; 11q13.2 - q13.3;<br>13q13.3; 13q14.11 - q14.13;<br>13q22.1; 14q11.2 - q21.2; 14q13.1 - q13.2; 14q23.2 - q23.3; 15q11.1 - q11.2; 15q21.2; 16q22.1; 17q12 - q21.32; 18q12.1;<br>21q22.12 - q22.13; 22q11.1 - q13.33; 22q12.1 - q12.2; 22q12.2 - q12.3; 22q13.1; 22q13.1 - q13.2; Xp22.33; Xq12 |
| **Recurrent gains** | 7 p15.2; 15q11.1-q11.2* | 1q21.1-q25.1 | 1p36.33 - p11.2; 1p31.1; 1q21.1 - q44; 2q31.1; 3q27.1; 4p16.1;<br>4q31.21; 5p12; 7p22.3 - p11.2; 7p15.2; 7q11.21 - q36.3; 8p11.22;<br>8q24.3; 10p12.2; 10q26.3; 11p15.5; 11q13.1; 14q11.2;<br>14q21.1; 14q32.2; 14q32.33; 15q11.1 - q11.2; 15q22.31; 16p13.3;<br>16q22.1; 17p13.1; 17q11.1 - q25.3; 17q21.32; 17q25.3;<br>19p13.3; 19p13.2; 19p13.12; 19q11 - q13.43; 19q13.11;<br>19q13.42; 20p13 - p11.1; 20q11.21 - q13.33; 22q12.2; 22q13.2;<br>22q13.31 |
| **Recurrent deletions** | - | - | 1p31.1; 5p15.33**; 9p21.3; 12p13.31; 22q11.23 |
| **Recurrent amplifications** | - | - | 3q26.1***; 7p11.2 |

\*one sample showed gain while the other one loss of the same region; \*\* one sample showed deletion while the other one loss of the same region;
\*\*\* one sample showed amplification while the other one gain of the same region

Furthermore, pilocytic astrocytomas showed distinct changes that were found in grade I tumors but not seen in grades II, III and IV. Such exclusive changes comprised only losses and they are shown in Table 2.

**Table 2.**
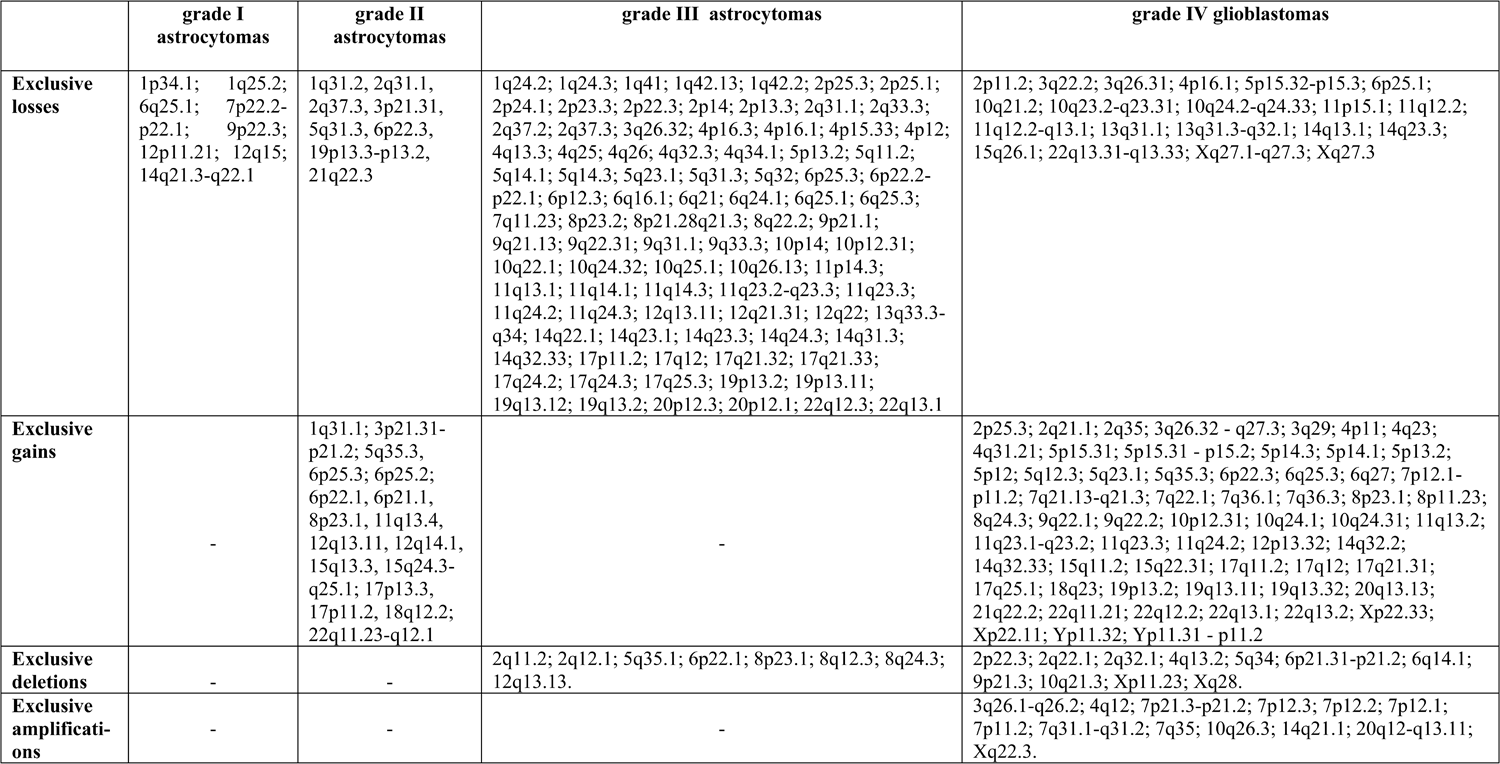
Exclusive changes found in different grades of astrocytomas.

|  | **grade I**<br>**astrocytomas** | **grade II**<br>**astrocytomas** | **grade III astrocytomas** | **grade IV glioblastomas** |
| --- | --- | --- | --- | --- |
| **Exclusive losses** | 1p34.1; 1q25.2;<br>6q25.1; 7p22.2-p22.1; 9p22.3;<br>12p11.21; 12q15;<br>14q21.3-q22.1 | 1q31.2, 2q31.1,<br>2q37.3, 3p21.31,<br>5q31.3, 6p22.3,<br>19p13.3-p13.2,<br>21q22.3 | 1q24.2; 1q24.3; 1q41; 1q42.13; 1q42.2; 2p25.3; 2p25.1;<br>2p24.1; 2p23.3; 2p22.3; 2p14; 2p13.3; 2q31.1; 2q33.3;<br>2q37.2; 2q37.3; 3q26.32; 4p16.3; 4p16.1; 4p15.33; 4p12;<br>4q13.3; 4q25; 4q26; 4q32.3; 4q34.1; 5p13.2; 5q11.2;<br>5q14.1; 5q14.3; 5q23.1; 5q31.3; 5q32; 6p25.3; 6p22.2-p22.1; 6p12.3; 6q16.1; 6q21; 6q24.1; 6q25.1; 6q25.3;<br>7q11.23; 8p23.2; 8p21.28q21.3; 8q22.2; 9p21.1;<br>9q21.13; 9q22.31; 9q31.1; 9q33.3; 10p14; 10p12.31;<br>10q22.1; 10q24.32; 10q25.1; 10q26.13; 11p14.3;<br>11q13.1; 11q14.1; 11q14.3; 11q23.2-q23.3; 11q23.3;<br>11q24.2; 11q24.3; 12q13.11; 12q21.31; 12q22; 13q33.3-q34; 14q22.1; 14q23.1; 14q23.3; 14q24.3; 14q31.3;<br>14q32.33; 17p11.2; 17q12; 17q21.32; 17q21.33;<br>17q24.2; 17q24.3; 17q25.3; 19p13.2; 19p13.11;<br>19q13.12; 19q13.2; 20p12.3; 20p12.1; 22q12.3; 22q13.1 | 2p11.2; 3q22.2; 3q26.31; 4p16.1; 5p15.32-p15.3; 6p25.1;<br>10q21.2; 10q23.2-q23.31; 10q24.2-q24.33; 11p15.1; 11q12.2;<br>11q12.2-q13.1; 13q31.1; 13q31.3-q32.1; 14q13.1; 14q23.3;<br>15q26.1; 22q13.31-q13.33; Xq27.1-q27.3; Xq27.3 |
| **Exclusive gains** | - | 1q31.1; 3p21.31-p21.2; 5q35.3,<br>6p25.3; 6p25.2;<br>6p22.1, 6p21.1,<br>8p23.1, 11q13.4,<br>12q13.11, 12q14.1,<br>15q13.3, 15q24.3-q25.1; 17p13.3,<br>17p11.2, 18q12.2;<br>22q11.23-q12.1 | - | 2p25.3; 2q21.1; 2q35; 3q26.32 - q27.3; 3q29; 4p11; 4q23;<br>4q31.21; 5p15.31; 5p15.31 - p15.2; 5p14.3; 5p14.1; 5p13.2;<br>5p12; 5q12.3; 5q23.1; 5q35.3; 6p22.3; 6q25.3; 6q27; 7p12.1-p11.2; 7q21.13-q21.3; 7q22.1; 7q36.1; 7q36.3; 8p23.1; 8p11.23;<br>8q24.3; 9q22.1; 9q22.2; 10p12.31; 10q24.1; 10q24.31; 11q13.2;<br>11q23.1-q23.2; 11q23.3; 11q24.2; 12p13.32; 14q32.2;<br>14q32.33; 15q11.2; 15q22.31; 17q11.2; 17q12; 17q21.31;<br>17q25.1; 18q23; 19p13.2; 19q13.11; 19q13.32; 20q13.13;<br>21q22.2; 22q11.21; 22q12.2; 22q13.1; 22q13.2; Xp22.33;<br>Xp22.11; Yp11.32; Yp11.31 - p11.2 |
| **Exclusive deletions** | - | - | 2q11.2; 2q12.1; 5q35.1; 6p22.1; 8p23.1; 8q12.3; 8q24.3;<br>12q13.13. | 2p22.3; 2q22.1; 2q32.1; 4q13.2; 5q34; 6p21.31-p21.2; 6q14.1;<br>9p21.3; 10q21.3; Xp11.23; Xq28. |
| **Exclusive amplifications** | - | - | - | 3q26.1-q26.2; 4q12; 7p21.3-p21.2; 7p12.3; 7p12.2; 7p12.1;<br>7p11.2; 7q31.1-q31.2; 7q35; 10q26.3; 14q21.1; 20q12-q13.11;<br>Xq22.3. |

### CNA in diffuse astrocytomas (grade II)

When analyzing joint CNA for diffuse astrocytomas (grade II) we also found that samples shared specific common variations. It is obvious from the Table 1 that the aberrant regions shared between grade II astrocytomas were less abundant than the number of aberrations found to be shared between grade I. The changes commonly found in grade II tumors included losses on chromosome cytobands 1p36.33-p11.2 and 1q21.1 and gains on 1q21.1-q25.1. Changes shared by grade II tumors were not found (repeated) in any of the grade I cases (Table 1).

We also sought for CNA aberrations exclusive for grade II tumors and found losses as well as gains as shown in Table 2.

Next we decided to investigate whether any specific affected region appearing in any (either or whichever) grade I patient could also be found in any given grade II patient. By such approach we discovered that regions on chromosomes 17 and 19 were characteristic for low grade astrocytomas since they were shared by at least two low grade patients. More precisely, patients suffering from astrocytoma grade I or II shared losses on region 19q13.11-q13.43. On region 17q21.2-q21.31 one grade I tumor showed loss, while grade II gain.

### CNA in anaplastic astrocytomas (grade III) and glioblastoma (grade IV)

Unlike recurrent changes found in grade II tumors, the observed concurrent changes in grades III and IV tumors were numerous. There were altogether 127 CNA that could be found across grades III and IV astrocytomas. They are listed in Table 1. It is interesting that both regions found to be affected in grade II astrocytomas, 1p36.33- p11.2 and 1q21.1, were also repeatedly affected in high grade tumors (Table 1).

CNA that were concurrent for pilocytic (grade I), anaplastic (grade III) and glioblastoma (grade IV) cases were: losses on 3q26.2; 4q28.2; 5q23.2; 6q13; 7p15.2 (gain and loss both); 10q11.21-q11.22; 10q21.3-q22.1; 11p15.4; 12p13.2 (deletion in high grades loss in pilocytic); 14q11.2; 14q13.1-q13.2; 15q11.1-q11.2; 18p11.22. Gains that were shared among I, III and IV grades were 7p15.2 and 15q11.1-q11.2. Those concurrent changes may indicate early events, since they were found both in low and high astrocytoma grades. Interestingly, region 17q22-q23.1 that lies within 17q11.1-25.3 was lost in grade I tumors, while the larger region 17q11.1-25.3 was gained in tumors with grades III and IV.

We noticed that grade III had an extensive number of exclusive aberrations consisting solely of losses and deletions without any specific gain or amplification. Exclusive grade III losses and deletions are shown in Table 2.

Glioblastomas (grade IV) also showed an extensive number of exclusive aberrations, even higher than the number found in other grades. Such unique CNA were losses and deletions, but also gains and amplifications and are listed in Table 2.

### Assigning the most frequently aberrant regions

We were also interested in how often specific regions were concurrent in our total astrocytoma patients. Therefore we searched for most frequently aberrant regions shared among the highest number of investigated patients. We defined the region as frequent when the same CNV was detected in 3 or more patients.

Four patients shared losses on 3p26.3-p12.3; 3q26.1; 10q11.21-q26.3; 13q14.11- q14.13; 22 q13.1; five patients shared losses on 9p24.3-p11.2; 9p21.3; 9p21.3-p13.3; 10p15.3- p11.1; 12p13.31; 22q11.23; and seven patients shared losses on 14q11.2.

Gains that 3 or more patients had in common were as follows: four patients shared gains on chromosome 2q31.1; 4p16.1; 7p15.2; 8q24.3; 11p15.5; 17p13.1; 17q25.3; 19p13.2; 19q11-q13.43; 19q13.42; 20q11.21-q13.33; five patients shared gains on 7q11.21-q36.3; 7p22.3-p11.2; 8p11.22; 10p12.2; 14q11.2; 15q11.1-q11.2. We presumed that those regions could harbor genes important for glial tumorigenesis. The regions with highest frequencies are shown in Fig 1.

**Fig 1.**
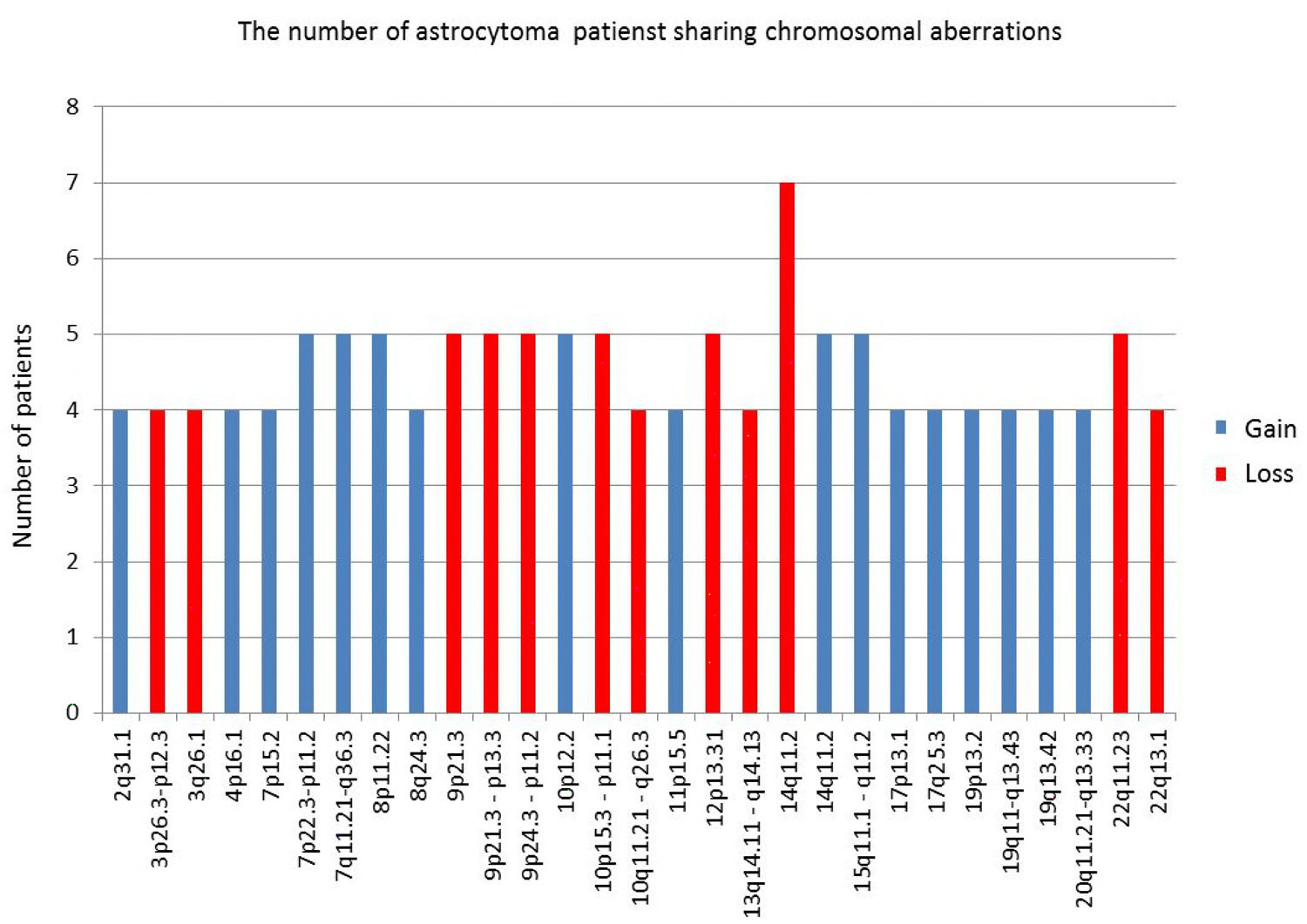
Frequent aberrant regions shared among the highest number of investigated patients.

### Broad regions

It has been shown that gliomas can display both focal and broad aberrations of their genome (Beroukhim et al, 2007). Therefore, we also decided to assess the regions showing broad changes. The analysis was performed by the following criteria: minimum log ratio for gains is 0.25 and for losses -0.25, minimal size of CNAs 2Mb and minimum number of consecutive probes 3. The list of broad aberrations found in our investigated group of astrocytomas is shown in Table 3.

**Table 3.** The regions showing broad changes

| chr | Regions | Change | WHO grade | References |
| --- | --- | --- | --- | --- |
| 1 | chr1p36.33-p11.2<br>chr1p36.33-p36.32<br>chr1p36.33-p35.2 | Loss | AII;<br>AIII;<br>2XG | Barbashina et al, 2005;<br>Beroukhim et al, 2007; Ichimura<br>et al, 2008; Yin et al, 2009 |
| 1 | chr1p36.33-p11.2 | Gain | 2XG |  |
| 1 | chr1q21.1-q25.1<br>chr1q21.1-q44 | Gain | AII; G | Mohapatra et al, 2011 |
| 3 | chr3p26.3-p12.3 | Loss | 4XG | Hesson et al, 2007; Mohapatra<br>et al, 2011 |
| 3 | chr3q26.32-q27.3<br>chr3q26.2-q29 | Gain | 2XG | Mohapatra et al, 2011 |
| 5 | chr5p15.31-p12<br>chr5p15.33-p11 | Gain | 2XG | Mohapatra et al, 2011;<br>Yang et al, 2011 |
| 7 | chr7 | Gain | 7XG | Beroukhim et al, 2007; Yin et<br>al, 2009 |
| 9 | chr9p24.3-p21.3 | Loss | 6XG | Yin et al, 2009;<br>Mohapatra et al, 2011 |
| 10 | chr10p15.3-p11.1 | Loss | 4XG | Mohapatra et al, 2011 |
| 10 | chr10q23.1-q26.3<br>chr10q11.21-q26.3<br>chr10q24.2-q24.33 | Loss | 4XG | Yin et al 2009;<br>Mohapatra et al, 2011 |
| 13 | chr13q21.2-q31.1<br>chr13q12.11-q31.3 | Loss | AIII; G | Yin et al, 2009;<br>Mohapatra et al, 2011 |
| 16 | chr16p13.3-p11.2<br>chr16p13.11-p12.1<br>chr16p12.3-p11.2 | Loss | AI; 2XG | Mohapatra et al, 2011 |
| 17 | chr17q11.1-q25.3 | Gain | 2XG | Yin et al 2009 |
| 19 | chr19p13.3-p12 | Gain | 4XG | Mohapatra et al, 2011 |
| 19 | chr19q11-q13.43 | Gain | 3XG | Barbashina et al, 2005;<br>Mohapatra et al, 2011 |
| 20 | chr20p13-p11.1 | Gain | 3XG | Brunner et al, 2000 |
| 20 | chr20q11.21-q13.33 | Gain | 3XG | Mohapatra et al, 2011 |
| 22 | chr22q11.21-q13.33 | Loss | 3XG | Mohapatra et al, 2011 |

Seven glioblastoma patients harbored amplification of chromosome 7 (trisomy of the whole chromosome 7) (Table 3). Broad region losses were at: chr1p36.33-p11.2; chr1p36.33- p36.32; chr1p36.33-p35.2; chr3p26.3-p12.3; chr9p24.3-p21.3; chr10p15.3-p11.1; chr10q23.1- q26.3; chr10q11.21-q26.3; chr10q24.2-q24.33; chr13q21.2-q31.1; chr13q12.11-q31.3; chr16p13.3-p11.2; chr16p13.11-p12.1; chr16p12.3-p11.2; chr22q11.21-q13.33 and broad regions gains at: chr1p36.33-p11.2; chr1q21.1-q25.1; chr1q21.1-q44; chr3q26.32-q27.3; chr3q26.2-q29; chr5p15.31-p12; chr5p15.33-p11; chr7; chr17q11.1-q25.3; chr19p13.3-p12; chr19q11-q13.43; chr20p13-p11.1; chr20q11.21-q13.33. Benign pilocytic astrocytomas lacked any of those types of changes.

### CNA in autologous blood sample DNA

Since aCGH uses as reference genome on a chip obtained from a pool of healthy individuals, we were interested whether some of the observed changes could be attributed to specific population polymorphisms. To ascertain whether some of the aberrations were of the constitutive nature, we also analyzed the autologous blood samples by comparing it to the reference DNA on the chip. The blood samples were from two patients, one suffering from pilocytic and one from glioblastoma. Neither of the two autologous blood samples harbored broad aberrations that are shown in Table 3. Autologous blood DNA from pilocytic astrocytoma sample showed altogether 23 copy number changes of which there were 3 amplifications, 8 gains, 9 losses and 3 deletions. The majority of changes (68%) from autologous constitutive DNA were repeated in the belonging tumor DNA (15/22) which may indicate individual or population CNV, but also an inborn susceptibility. Four amplifications and 3 deletions were exclusive for the blood DNA indicating probable population genetic variation.

Interestingly, all alterations noted for normal blood DNA from glioblastoma patient were repeated in the DNA of the belonging tumor, so there were no exclusive changes for autologous blood DNA of the tested glioblastoma patient. Shared alterations for tumor and blood DNA of this glioblastoma patient were 3 amplifications, 6 gains, 9 losses and 3 deletions.

### Functional Analysis by GISTIC2.0.23 Identified Significant Genomic Targets in Astrocytomas

With the objective to interpret and draw conclusions from our raw data and results we performed bioinformatics analyses. GISTIC identifies those regions of the genome that are aberrant more often than would be expected by chance. Greater weight is given to high amplitude gains or deletions that are less likely to represent random aberrations (Beroukhim et al, 2007). The GISTIC algorithm identifies likely somatic driver copy number CAN by evaluating the amplitude and frequency of either amplified or deleted observation (Mermel et al. 2011). To find statistically relevant recurrent CNA GISTIC determined focally amplified (red) and deleted (blue) regions plotted along the genome (Fig 2).

**Fig 2.**
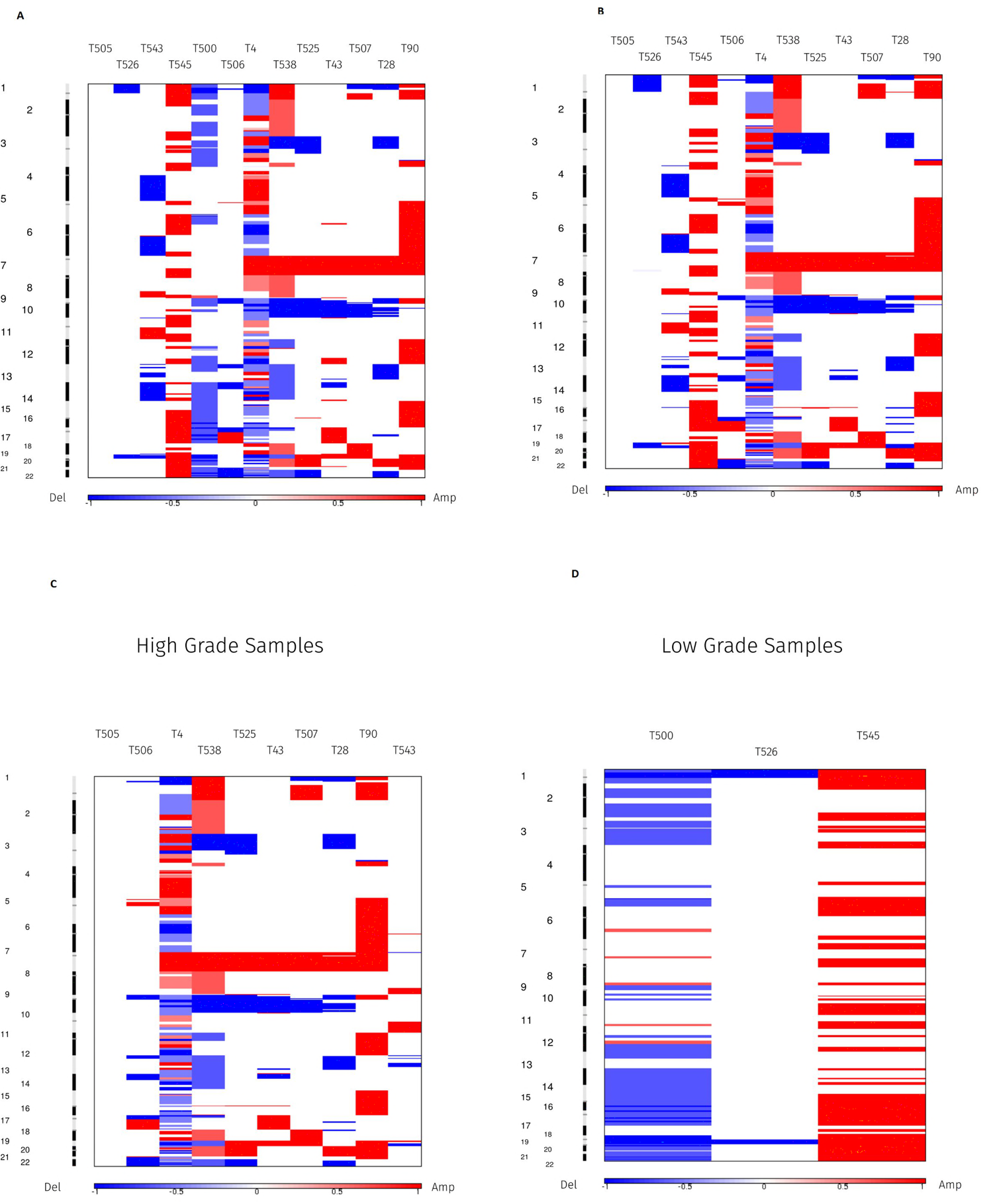
Heat map images of four different intercranial astrocytic brain tumor sample pools based on total segmented DNA copy number variation profiles. Images were analysed using GISTIC (v2.0.23). In each heat map, the samples are arranged from left to right, and chromosome arrangement flows vertical, top to bottom ordering. Red represents CN gain and blue represents CN loss.

Furthermore, GISTIC2.0.23 was also used to identify significant amplification and deletion events assigned to malignancy grades. To this end, we computed GISTIC CNA amplification/deletion plots, segmented CN heat plots and identify genes within those regions. Moreover, we utilized a range of cutoff values in the analysis to identify segments of significance.

### Results on total sample analysis

The results of GISTIC algorithm analysis identified regions of aberration that are more likely to drive cancer pathogenesis. A number of regions of recurrent CN gains and losses have been identified across all samples. Their genomic locations including the number of the associated genes as a function of the corresponding q-value cutoff criterion is summarized in Table 4. For technical reasons we had to exclude one pilocytic astrocytoma case from bioinformatics analysis.

**Table 4.** List of regions and their associated genes located in the most common sections of recurrent DNA CNAs, derived from analysis conducted on our total astrocytic brain tumor sample.

| Focal event | Cytoband | q-value | Genomic position (hg19) | 0,45 | 0,35 | 0,25 | 0,15 | 0,05 |
| --- | --- | --- | --- | --- | --- | --- | --- | --- |
| Amplification | 3q28 | 0,36509 | chr3:169432744-198022430 | 217 | 0 | 0 | 0 | 0 |
| Amplification | 4p16.3 | 0,36509 | chr4:1309269-1885110 | 10 | 0 | 0 | 0 | 0 |
| Amplification | 8q24.3 | 0,36509 | chr8:144884270-146364022 | 62 | 0 | 0 | 0 | 0 |
| Amplification | 12q13.3 | 0,36509 | chr12:57863606-58162220 | 20 | 0 | 0 | 0 | 0 |
| Amplification | 12p13.32 | 0,36509 | chr12:3736374-4718832 | 9 | 0 | 0 | 0 | 0 |
| Amplification | 14q32.33 | 0,36509 | chr14:106400482-107349540 | 3 | 0 | 0 | 0 | 0 |
| Amplification | 17q25.3 | 0,36509 | chr17:78847437-79535591 | 25 | 0 | 0 | 0 | 0 |
| Amplification | 21q22.3 | 0,36509 | chr21:46788100-46974713 | 3 | 0 | 0 | 0 | 0 |
| Amplification | 22q13.33 | 0,36509 | chr22:50033682-51304566 | 40 | 0 | 0 | 0 | 0 |
| Total: |  |  |  | 389 | 0 | 0 | 0 | 0 |
| Deletion | 17p13.2 | 0,0027597 | chr17:1-7172830 | 176 | 176 | 176 | 176 | 213 |
| Deletion | 9p21.3 | 0,024206 | chr9:21030772-22655576 | 27 | 27 | 27 | 34 | 34 |
| Deletion | 13q12.11 | 0,024206 | chr13:19891479-20000549 | 1 | 1 | 1 | 1 | 1 |
| Deletion | 22q12.3 | 0,037797 | chr22:28350866-44221419 | 261 | 261 | 261 | 261 | 261 |
| Deletion | 10q24.2 | 0,059906 | chr10:93786297-105364688 | 171 | 171 | 171 | 178 | 0 |
| Deletion | 15q15.1 | 0,059906 | chr15:40736349-42074646 | 33 | 33 | 33 | 33 | 0 |
| Deletion | 16q22.1 | 0,059906 | chr16:66967463-74334046 | 130 | 130 | 130 | 138 | 0 |
| Deletion | 17q21.31 | 0,059906 | chr17:36996467-45200422 | 255 | 255 | 255 | 255 | 0 |
| Deletion | 20q11.22 | 0,059906 | chr20:6031411-36620782 | 240 | 240 | 240 | 240 | 0 |
| Deletion | 14q12 | 0,15415 | chr14:1-47314894 | 232 | 235 | 235 | 0 | 0 |
| Deletion | 3p14.3 | 0,15415 | chr3:57198280-58186818 | 9 | 9 | 9 | 0 | 0 |
| Deletion | 5q23.2 | 0,15415 | chr5:125909127-126208712 | 3 | 3 | 3 | 0 | 0 |
| Deletion | 12q21.33 | 0,15415 | chr12:31143267-133851895 | 879 | 879 | 879 | 0 | 0 |
| Deletion | 11p15.4 | 0,2165 | chr11:9277720-9686181 | 6 | 6 | 6 | 0 | 0 |
| Total: |  |  |  | 2423 | 2426 | 2426 | 1316 | 509 |

For exploratory reasons and with conscious danger of over-interpreting our results by including anecdotal structural changes, we decided to investigate CNA by lowering the cutoff of q-values. We utilized a range of cutoff values in the analysis to identify segments of significance. Such more permissive approach revealed some additional relevant regions. The results are presented in Table 4.

By focusing on 0.25 (q-value) criterion, no significant amplifications were detected, while at the same threshold level, significant deletions affected 14 chromosomal regions, out of which 4 (17p13.2, 9p21.3, 13q12.11, 22q12.3) remained significant even at 0.05 q-value cutoff level, all with GISTIC scores higher than 55.

### Results of malignant astrocytoma analysis (grade I pilocytic cases excluded)

Computational analysis using GISTIC was repeated as previously described. However, this reanalysis excluded benign astrocytoma cases. Such approach resulted in three novel, previously disregarded regions at q-value threshold of 0.25 to be identified as significantly amplified (3q28, 14q32.33, 18q12.2), while the number of significantly deleted regions decreased by more than a half (from 14 to 6). Two of those (17p13.2, 9p21.3) still remained significant at threshold level of q-value=0.05. Out of 6 remaining regions 4 overlapped with those in the previous analysis (17p13.2, 13q12.11; 10q24.2, 9p21.3). The region 14q32.33 found amplified on the total sample at q-0.45 was amplified in malignant cases at q-0.25. Table 5 and Fig 2B summarize the obtained result.

**Table 5.** List of regions and their associated genes located in the most common sections of recurrent DNA CNAs, derived from analysis conducted on malignant astrocytic brain tumor samples. Benign cases were excluded

| Focal event | Cytoband | q-value | Genomic position (hg19) | Gene Count ( q-value cutoff ) |  |  |  |  |
| --- | --- | --- | --- | --- | --- | --- | --- | --- |
|  |  |  |  | 0,45 | 0,35 | 0,25 | 0,15 | 0,05 |
| Amplification | 3q28 | 0,18103 | chr3:169432744-198022430 | 217 | 217 | 217 | 0 | 0 |
| Amplification | 14q32.33 | 0,18103 | chr14:106400482-107349540 | 3 | 3 | 3 | 0 | 0 |
| Amplification | 18q12.2 | 0,18103COG, | chr18:34934984-35256171 | 3 | 3 | 3 | 0 | 0 |
| Amplification | 1p36.32 | 0,26407 | chr1:2215776-2801808 | 14 | 14 | 0 | 0 | 0 |
| Amplification | 4p16.3 | 0,26407 | chr4:1309269-1885110 | 10 | 10 | 0 | 0 | 0 |
| Amplification | 8q24.3 | 0,26407 | chr8:144884270-146364022 | 62 | 62 | 0 | 0 | 0 |
| Amplification | 12q13.3 | 0,26407 | chr12:57863606-58162220 | 20 | 20 | 0 | 0 | 0 |
| Amplification | 12p13.32 | 0,26407 | chr12:3736374-4718832 | 9 | 9 | 0 | 0 | 0 |
| Amplification | 20q13.33 | 0,26407 | chr20:61784176-63025520 | 55 | 55 | 0 | 0 | 0 |
| Amplification | 21q22.3 | 0,26407 | chr21:46788100-46976402 | 3 | 3 | 0 | 0 | 0 |
| Amplification | 22q13.33 | 0,26407 | chr22:50033682-51304566 | 40 | 40 | 0 | 0 | 0 |
| Total: |  |  |  | 436 | 436 | 223 | 0 | 0 |
| Deletion | 17p13.2 | 0,0047853 | chr17:1-7172830 | 176 | 176 | 176 | 176 | 176 |
| Deletion | 9p21.3 | 0,0068152 | chr9:20621756-25684739 | 34 | 34 | 34 | 34 | 34 |
| Deletion | 10q24.2 | 0,058622 | chr10:93783434-105364688 | 171 | 171 | 171 | 178 | 0 |
| Deletion | 14q21.3 | 0,058622 | chr14:48262357-51191843 | 20 | 20 | 20 | 20 | 0 |
| Deletion | 1p36.11 | 0,23541 | chr1:24991120-31189272 | 97 | 97 | 97 | 0 | 0 |
| Deletion | 13q12.2 | 0,23541 | chr13:1-38115337 | 119 | 119 | 119 | 0 | 0 |
| Total: |  |  |  | 617 | 617 | 617 | 408 | 210 |

### The results of high grade samples analysis (grades III and IV)

By excluding low grade samples (grades I and II) and repeating the analysis on high grade samples including the most aggressive type glioblastoma (grade IV), none of the previously identified amplified regions, classified as statistically significant at 0.25 q-value threshold, were observed. On the other hand, all previously identified deletions on malignant group were still present constituting a stable result when it comes to the identified deletion events (Table 6, Fig 2C). By increasing the cutoff value from 0.25 to 0.35, significant amplifications become evident and in line with three of the previously identified ones, 3q28; 12q13.3 and 21q22.3. One of which (3q28) was significant at 0.25 threshold for malignant astrocytoma group and at p-0.45 threshold level on our total analyzed sample, thus in line with the assumed, stable cross sample identified amplification region.

**Table 6.** List of regions and their associated genes located in the most common sections of recurrent DNA CNAs, derived from analysis conducted involving high grade samples (grades III and IV).

| Focal event | Cytoband | q-value | Genomic position (hg19) | Gene Count ( q-value cutoff ) |  |  |  |  |
| --- | --- | --- | --- | --- | --- | --- | --- | --- |
|  |  |  |  | 0,45 | 0,35 | 0,25 | 0,15 | 0,05 |
| Amplification | 3q28 | 0,29658 | chr3:169432744-198022430 | 217 | 217 | 0 | 0 | 0 |
| Amplification | 12q13.3 | 0,29658 | chr12:57863606-58210057 | 24 | 24 | 0 | 0 | 0 |
| Amplification | 21q22.3 | 0,29658 | chr21:46788100-46974713 | 3 | 3 | 0 | 0 | 0 |
| Total: |  |  |  | 244 | 244 | 0 | 0 | 0 |
| Deletion | 9p21.3 | 0,049849 | chr9:20621756-25684739 | 34 | 34 | 34 | 34 | 34 |
| Deletion | 17p13.2 | 0,05166 | chr17:1-7172830 | 176 | 176 | 176 | 176 | 0 |
| Deletion | 10q24.2 | 0,051797 | chr10:93783434-105367017 | 171 | 171 | 171 | 178 | 0 |
| Deletion | 14q21.3 | 0,051797 | chr14:48262357-51191843 | 20 | 20 | 20 | 20 | 0 |
| Deletion | 1p36.11 | 0,21828 | chr1:24995465-31189272 | 97 | 97 | 97 | 0 | 0 |
| Deletion | 13q12.11 | 0,21828 | chr13:21324433-21947432 | 5 | 5 | 5 | 0 | 0 |
| Total: |  |  |  | 503 | 503 | 503 | 408 | 34 |

### The results of low grade samples analysis (grades I and II)

The last computation in this GISTIC utility series was the analysis involving low grade samples (AI and AII). Table 7 and Fig 2D contain the obtained result. The lack of significant deletions, as well as of amplifications was evident at q-value of 0.25.

**Table 7.** List of regions and their associated genes located in the most common sections of recurrent DNA CNAs, derived from analysis conducted involving intracranial astrocytic brain tumor low grade samples (grades I and II).

| Focal event | Cytoband | q-value | Genomic position (hg19) | Gene Count ( q-value cutoff ) |  |  |  |  |
| --- | --- | --- | --- | --- | --- | --- | --- | --- |
|  |  |  |  | 0,45 | 0,35 | 0,25 | 0,15 | 0,05 |
| Deletion | 3p14.3 | 0,44232 | chr3:57198280-58187545 | 9 | 0 | 0 | 0 | 0 |
| Deletion | 5q21.2 | 0,44232 | chr5:1-180915260 | 1050 | 0 | 0 | 0 | 0 |
| Deletion | 11p15.4 | 0,44232 | chr11:1-135006516 | 1478 | 0 | 0 | 0 | 0 |
| Deletion | 13q12.2 | 0,44232 | chr13:1-38110402 | 119 | 0 | 0 | 0 | 0 |
| Deletion | 15q15.1 | 0,44232 | chr15:40735035-42074637 | 33 | 0 | 0 | 0 | 0 |
| Deletion | 16q22.2 | 0,44232 | chr16:66967005-75040923 | 138 | 0 | 0 | 0 | 0 |
| Deletion | 17q12 | 0,44232 | chr17:1-64211652 | 1112 | 0 | 0 | 0 | 0 |
| Deletion | 18q21.1 | 0,44232 | chr18:43535382-43924106 | 4 | 0 | 0 | 0 | 0 |
| Deletion | 20q11.22 | 0,44232 | chr20:6027505-36613689 | 240 | 0 | 0 | 0 | 0 |
| Deletion | 22q12.3 | 0,44232 | chr22:28346538-44221618 | 261 | 0 | 0 | 0 | 0 |
| Total: |  |  |  | 4444 | 0 | 0 | 0 | 0 |

To sum up the results of GISTIC 2.0.23 analysis of CN profiles above, we can point out that regions identified as significantly deleted in high grade samples were: 9p21.3; 17p13.2; 10q24.2; while regions 14q21.3; 1p36.11 and 13q12.11 surfaced only in malignant cases indicating their involvement as later events.

Regions significantly amplified and connected to pronounced malignancy were 3q28; 12q13.3 and 21q22.3, of which last two emerged only in high grade cases, while 3q28 was constantly found. None of the above aberrations were significant in low grade astrocytoma tumors. Regions 17q25.3 and 8q24.3 that were significantly amplified on our total sample did not emerge in subsequent analyses and therefore may be characteristic for lower grade astrocytomas. Of note is that deletions 3p14.3; 11p15.4; 15q15.1; 16q22.1; 20q11.22 and 22q12.3 were all found in low grade samples at threshold level of q-0.45 and also on our total sample at q-0.25, but were not repeatedly found in high grades. These findings indicate that these regions and genes within may also be involved as early events.

### Computing GISTIC Heat Maps for the previous analyses

Chromosomal alterations based on DNA CN changes in all four case studies are illustrated using heat maps (Fig 2). Upon visual inspection clear distinction between low and high grade samples is evident, with high grade samples closely reflecting the general heat map images of the entire batch (Fig 2C and Fig 2D). Moreover, an almost systematic amplification of segments across majority of samples in chromosome 7 and respective deletion in chromosome 10 could be observed.

### Assessing Functional Features of Genes Identified by GISTIC (Relevant annotated genes)

The most significant amplifications and deletions identified by GISTIC as previously described revealed several aberrant regions and genes within which were further investigated using functional enrichment strategies as implemented in DAVID. Out of 840 CNA associated genes, according to DAVID, 81 were not linked to any known pathway or function based annotation category. Of the remaining 759 associated, only 65 genes assigned to a pathway or a functional category were significantly over-represented (Bonforroni and BH adjusted p-value0.05) via gene annotation within the identified CNAs. Fig 3 summarizes the distribution of those 65 identified genes across different enrichment categories as well as the information regarding their shared contribution to each of the indicated individual categories. The list of the associated 65 genes is located in Table 8 while in EV1 Table their distribution across cytobands with corresponding significance metrics can be found.

**Fig 3.**
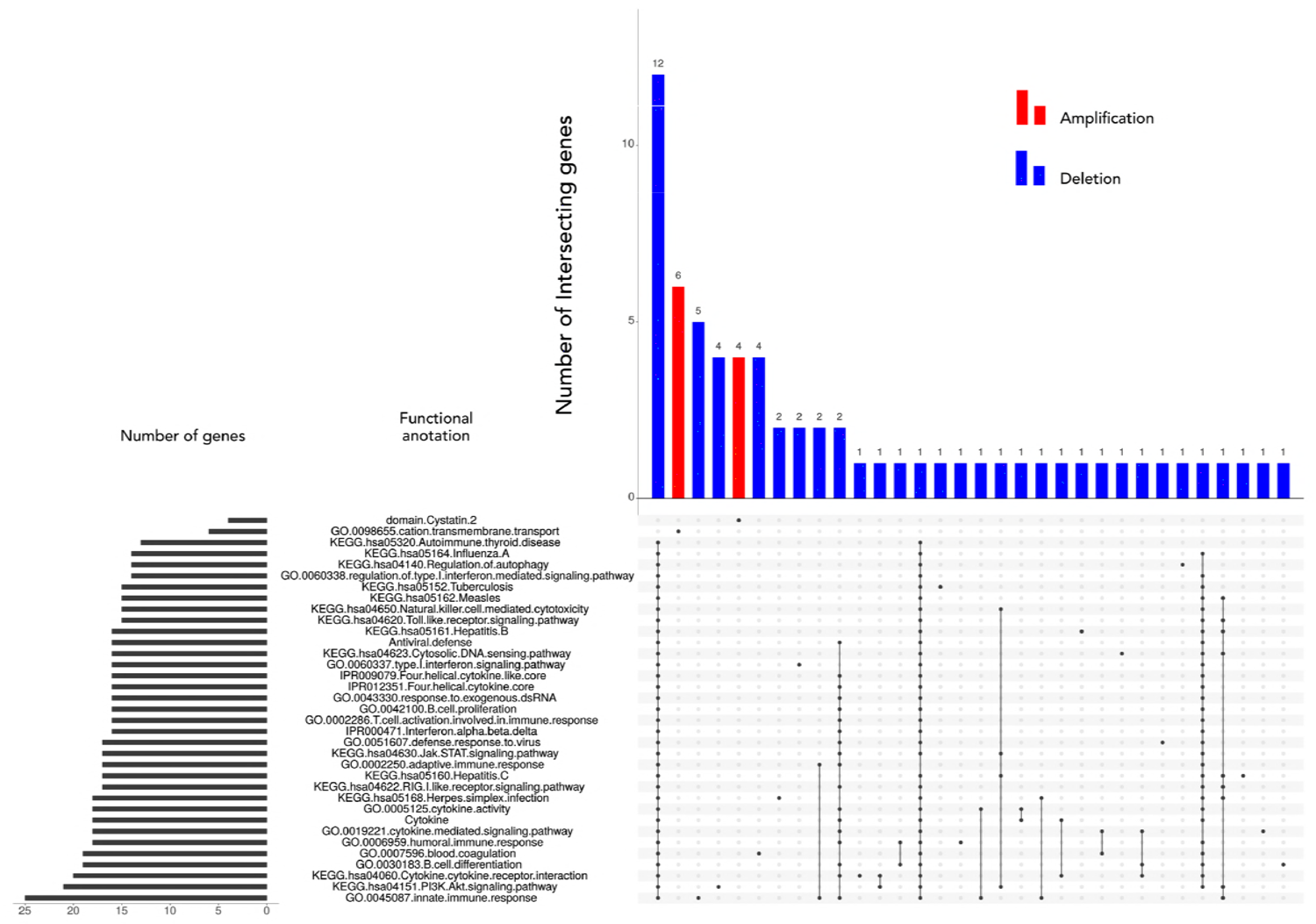
Matrix layout for all 65 genes across 35 functional categories (association calculated by DAVID), sorted by size. Dark circles in the matrix indicate functional categories with genes that are part of the intersecting groups, that is, are associated with each category of the set. Bar plot above matrix depicts the number of shared genes, while the horizontal bar plot on the left reflects the number of genes within each group. Blue and red colored bars indicate the respective aberration.

**Table 8.** List of genes within CNAs associated with significantly enriched functional categories as calculated using DAVID computational strategy (Bonforroni and BH adjusted p-value threshold se to α = 0.05).

| Gene ID |
| --- |
| AHSG, ATP13A3, ATP13A4, ATP13A5, BLNK, C1QBP, CCNA1, CHUK, CLDN7, CLEC10A, CXCL16, ENTPD1, FCN3, FETUB, FGF8, FGF9, FGR, FLT1, FLT3, GABARAP, GP1BA, HHEX, HMGB1, HPS6, HRG, HTR3C, HTR3D, HTR3E, IFI6, IFNA1, IFNA10, IFNA13, IFNA14, IFNA16, IFNA17, IFNA2, IFNA21, IFNA4, IFNA5, IFNA6, IFNA7, IFNA8, IFNB1, IFNE, IFNW1, IL17D, KNG1, NFKB2, NLRP1, P2RX1, P2RX5, PIK3AP1, POLR1D, RFXAP, RTN4RL1, SMPDL3B, SOS2, SRSF4, TAF5, TNFRSF19, TRIM8, XAF1, YTHDF2, YWHAE, ZNF683 |

In order to position functionally relevant over-represented genes, we allocated them to the regions that GISTIC 2.0.23 identified as significantly deleted in high grade samples. In such a manner region 14q21.3 comprised gene SOS2, region 1p36.11 comprised genes FCN3, ZNF683, region 13q12.11 genes FGF9, FLT1, FLT3, HMGB1, IL17D, POLR1D, TNFRSF19. Region 9p21.3 comprises genes for several interferon molecules IFNA1, IFNA10, IFNA13, IFNA14, IFNA16, IFNA17, IFNA2, IFNA4, IFNA5, IFNA6, IFNA7, IFNA8, IFNB1; while at 17p13.2(1) resided genes C1QBP, CLDN7, CLEC10A, CXCL16, DHX33, GABARAP, GP1BA, NLRP1, P2RX1, P2RX5, RTN4RL1, XAF1, YWHAE and DVL2. At 10q24 genes BLNK, CHUK, ENTPD1, FGF8, HPS6, NFKB2, PIK3AP1, TAF5, TRIM8 and WNT8B were allocated.

The significantly amplified region in high grades 3q28 was the only one to harbor functionally relevant annotated gene - CLDN1.

The region suspected as an early event found in low grade tumor, 20q11.22, was the only region to harbor functionally relevant annotated gene - MAP1LC3A (LC3).

Annotated genes were comprised to other different regions as well. Region 1p35.3 harbored genes FGR, IFI6, SMPDL3B, SRSF4, YTHDF2, region 1q42.12 gene PARP1; region 3q27.3 genes AHSG, FETUB, HRG, HTR3C, HTR3D, KNG1, region 3q27.1 genes HTR3E and DVL3, region 3q29 genes ATP13A3, ATP13A4, ATP13A5, region 10q23.3 gene HHEX, region 13q13.3 genes CCNA1, RFXAP; while allocated at 4q27 was gene CCNA2 (cyclin A), at 5q23.1 gene LOX, at 5q32 gene CSNK1A1 and at 6q27 gene TFIID.

### Signaling pathways involved

To further elevate specificity of our analysis we restricted our functional pathway analysis to that contained within KEGG database preforming the enrichment analyses with p<0.05 as a cutoff criterion. As a result, only genes associated to deleted segments were significantly enriched in 18 out of 325 total Homo sapiens associated KEGG pathways.

Fig 4 illustrates this result indicating PI3K-Akt signaling pathway, Cytokine-cytokine receptor interaction, NOD-like receptor signaling pathways and pathways involved in HPV and herpes simplex infection being the most significantly represented in terms of the number of genes identified by GISTIC. Among the signaling pathways significant enrichments were detected in PI3K-Akt, Cytokine-cytokine receptor interaction, NOD-like receptor, Jak-STAT, RIG-II-like receptor and Toll-like receptor pathway. Also, pathways involved in HPV, herpes simplex, hepatitis C and B infections as well as Kaposi’s sarcoma-associated herpesvirus infection, tuberculosis, measles, influenza A were all significantly enriched too. Several pathways involved in inflammation – necroptosis, Cytosolic DNA-sensing pathway, Natural killer cell mediated cytotoxicity. And finally the pathway connected to autoimmune thyroid disease showed to be significant, too.

**Fig 4.**
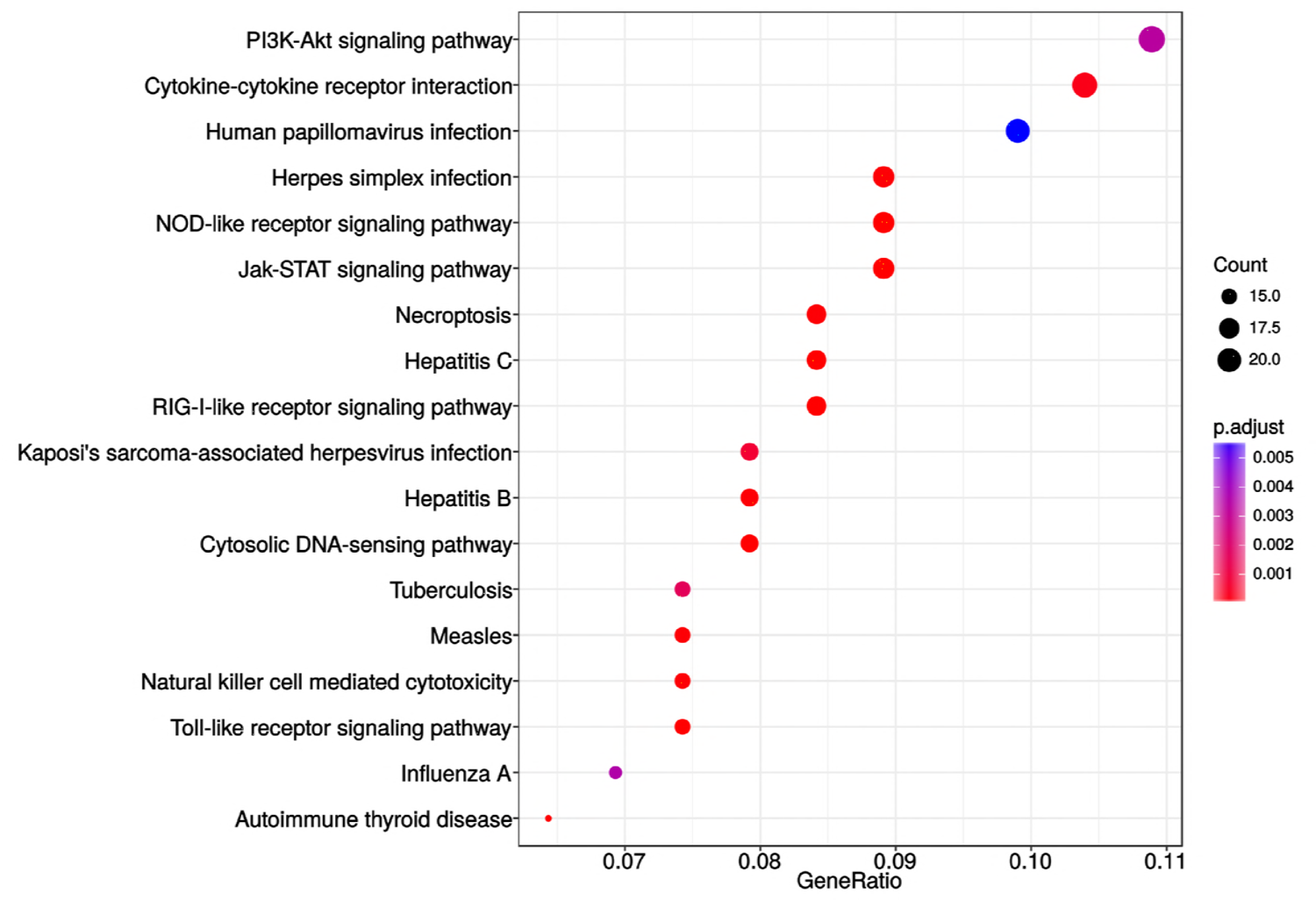
Enrichment analysis utilizing KEGG pathway database. Analysis included genes from all malignant samples associated to both deleted and amplified regions. No significant enrichment was associated to amplified segments.

To further investigate the role of these genes we plotted the enrichment map (EV Fig 1) revealing roughly equal systematic involvement of all genes across the identified pathways.

To see how many of the identified genes and across how many KEGG pathways underlined the enriched pathways, we preformed the set intersection analysis (EV Fig 2). The obtained result confirms previous indications asserting 12 out of 44 KEGG associated genes to be shared among 18 significantly enriched pathways.

Finally, we extracted the most important identified KEGG pathways shown in EV Fig 3. and labeled genes associated with deleted chromosomal regions.

## Discussion

In the present investigation we wanted to elucidate which chromosomal regions and annotated genes are involved in the genesis and progression of astrocytic brain tumors. Cancer genomes suffer many structural changes (Chin et al, 2011) and CNAs have been commonly found in glioma (Mohapatra et al, 2013). However, CNAs differ in their frequency of recurrence even among the patients suffering from the same type of malady. Which specific CNAs are attributed as early events and which are responsible for progression still remains to be fully understood.

On our total sample we found that the number of losses significantly exceeded the number of observed gains and amplifications. This finding is not unusual since it has been reported as a general pattern in cancer (Beroukhim et al, 2010) that losses are more frequent than amplifications. Furthermore, we have found that the mean number of CNA is much higher in malignancy grades III and IV when compared to lower grades. Also, a great number of aberrant regions were recurring in grades III and IV.

Our study also revealed similarities and differences in aberrant chromosomal regions across astrocytoma grades. The CNV that were found to be shared among patients with benign grade I pilocytic astrocytomas indicated relatively different pattern than observed in malignant group. It has been postulated that pilocytic astrocytomas differ from other histopathological types as they are slow-growing and non-infiltrative. Although they usually exhibit a normal karyotype, ~32% display chromosomal abnormalities. Few genetic events were reported for pilocytic astrocytoma (Marko & Weil, 2012; Jones et al, 2013; Pećina-Šlaus et al, 2014b) including gains of chromosomes 5, 7 and 8 and less frequently gains of chromosomes 1, 3, 4, 6, 9–17, 19–22 and monosomies of chromosomes 7, 8, and 17. Chromosomal regions that have been reported to hold abnormalities include 1p, 2p, 4q-9q and 13q and losses on 1p, 9q, 12q and 19–22 (Jones et al, 2009; Ward et al, 2010; Marko and Weil, 2012). The situation found in our study is compatible to some of the aberrations reported previously, but also differed from the literature. We found losses in pilocytic astrocytomas of which: 3q; 10q; 11p; 12p; 14q; 15q and 18p have not previously been reported, while there were fewer gains found in our study, only on 7p15.2 and 15q11.1-q11.2.

Grade II astrocytomas harbored only few recurrent aberrations, namely losses on 1p36.33-p11.2 and 1q21.1 and gains on 1q21.1 - q25.1. None of them recurred in grade I tumors. However, losses recurrent for grade II astrocytomas were also repeatedly affected in higher grade tumors.

Malignant high grades tumors, III and IV, on the other hand, harboured numerous recurrent changes, which indicates the augmentation of aberrations as the disease progresses.

The majority of CNA that have been reported in the literature were also discovered and confirmed with our experiments (Brunner et al, 2000; Hesson et al, 2007). However, the frequencies differed as well as their previous assignments to specific grade. Seifert et al (2015) in their computational study revealed similarities and differences in gene expression levels between astrocytomas of all four WHO grades. The authors report that transcriptional alterations of individual signaling pathways typically increase with WHO grade of astrocytoma. The high number of copy number changes found to be increasing with the grade can also be indicative of the acquisition of genomic instability in glioblastoma, especially since deleted regions may harbor genes involved in mismatch DNA repair.

The most common amplification - the one on chromosome 7 (Ichimura et al, 2008; Brennan et al, 2013), was also frequently found in our investigated sample with 77.8% of tumors displaying this type of change. Another frequent event - deletions of chromosome 10 (Ichimura et al, 2008), has been discovered in 88.9% of our patients. This finding is in accordance with literature were loss of heterozygosity on chromosomal arm 10p is commonly reported for high-grade gliomas, usually concentrated in the region 10p14-p15 (Fleischer et al, 2011; Verhaak et al, 2010). Our study found losses of 10p11.1-p15.3 region in 77.8% of glioblastomas.

Brennan et al (2013) found higher frequencies of the most common amplification events reported for astrocytoma, on chromosomes 7 (EGFR/MET/CDK6), 12 (CDK4 and MDM2) and 4 (PDGFRA), than other investigators. However, these highly frequent events were not frequently targeted in our study. Only 22.2% of our cases showed gain on chr12, and 28.6% showed gains on 4p16.1.

Several of our results corroborate the findings of previous studies regarding relevant genes (Brennan et al (2013). The most common genomic aberrations seen in glioblastomas, with 93% of samples harboring chromosome 7 amplifications (EGFR/MET/CDK6) and chromosome 10 deletions, is the amplification of EGFR gene found in 95% of tumors. Homozygous deletion spanning the Ink4a/ARF locus (Bidinotto et al, 2016) was found in 95% of cases. Our study showed EGFR amplification to be targeted in 89% of glioblastomas and one astrocytoma grade II, totaling of 90% of cases with amplified EGFR gene.

In order to comprehend CNA events in astrocytomas of different pathohistological types and identify alterations that are biologically and functionally significant we used GISTIC algorithm. We were interested to differentiate founder events and subclonal drivers from passenger mutations (Vogelstein et al 2013; Abou-El-Ardat et al, 2017). The software was utilized previously in numerous cancer studies, including lung (Bass et al, 2009), colorectal carcinoma (Firestein et al, 2008) and melanoma (Lin et al, 2008) and has facilitated the identification of new significant cancer associated targets.

By exploring a range of cutoff q-values we identified additional segments of significance. Thus, significant deletions affecting 14 chromosomal regions were found, out of which deletions of 17p13.2, 9p21.3, 13q12.11 and 22q12.3 remained significant even at 0.05 q-value. Of importance is that locus 9p22.1-p21.3 encompasses the CDKN2A gene (p16INK4a/p14ARF/p15INK4b locus) known to be frequently deleted in gliomas (Yin et al., 2009). Furthermore, RB gene is residing in the region 13q and p53 gene in 17p13. The study by Yin et al (2009) found that the long arm of chromosome 13 was lost in nearly 33% to 50% of cases.

When excluding pilocytic cases, the GISTIC reanalysis resulted in three novel, previously disregarded regions to be identified as significantly amplified, 3p28, 14q32.33 and 18q12.2. Since the number of significantly deleted regions decreased by more than a half, it seems that deletions are characteristic of benign cases. Two of deleted regions, 17p13.2 and 9p21.3 still remained significant at threshold level of q-value 0.05. Out of 6 remaining regions 4 overlapped with those in the previous analysis (17p13.2, 13q12.11; 10q24.2, 9p21.3). We can assume that these regions could represent the first events to be changed in the consecutive steps of gliomagenesis. Of note is that the region 14q32.33 found amplified on total sample at q-0.45 was amplified in malignant cases at q-0.25.

In the analysis on high grade astrocytomas (III and IV), none of the previously identified amplified regions, classified as significant at 0.25 q-threshold, were observed. On the other hand, all previously identified deletions found in malignant group were present constituting a stable result when it comes to the identified deletion events. When the cutoff value was raised to 0.35, significant amplifications became evident and in line with three previously identified ones of which 3q28 was significant at 0.25 threshold for malignant astrocytoma group and at p-0.45 on our total sample. This is in line with the identified stable cross sample amplification region.

Low grade astrocytomas demonstrated the lack of both significant deletions and amplifications, suggesting a general pattern associated to these grades. Common genetic changes and tumor associated mutations found in higher grade gliomas, p53, PDGF, p16 (CDKN2A), IDH1 and IDH2 are rarely reported in pilocytic astrocytomas, which is consistent to our results that also indicate lack of focal abnormalities in loci where those genes reside.

When grouping our sample according to malignancy grade, the regions identified as significantly deleted in group with grades III and IV were: 9p21.3; 17p13.2; 10q24.2; 14q21.3; 1p36.11 and 13q12.11, and significantly amplified regions were 3q28; 12q13.3 and 21q22.3. None of the above aberrations were significant for low grade astrocytoma tumors, and we believe they might be associated to progression events.

Regions 17q25.3 and 8q24.3that were found to be amplified on our total sample did not emerge in subsequent analyses and therefore may be characteristic for lower grade astrocytomas. Of note is that deletions 3p14.3, 11p15.4, 15q15.1, 16q22.1, 20q11.22 and 22q12.3 were all found in low grade samples at threshold level of q-0.45 and also on our total sample at q-0.25, but were not repeatedly found in high grades. These findings indicate that these regions and genes within may also be involved as early events. Although many observed changes were similar to the literary reports, some were identified for the first time in our patients and associated to progression or as an early event.

We could not establish any differences between IDH1 mutant and WT tumors in regards of presence of listed CNAs.

Even though drawing conclusions is complicated perhaps because of the inherent heterogeneity of astrocytomas (Sottoriva et al, 2013) and complexity of cancer genomes *per se*, our bioinformatics results indicate compatibility with the previously reported regions. At first cancer-related aCGH studies have showed a high level of discordance in the reported genomic aberrations (Ali et al, 2017) guiding to conclusions that random mutations and CNA are prevalent. However, newly developed tool GISTIC can distinguish which CNA are more functionally relevant to the cancer evolution. The accordance rate among different studies improved and a concordant picture of biologically significant CNAs in the glioma genome emerged (Beroukhim et al, 2007).

Genes known to be the most frequently amplified in glioblastoma, EGFR, CDK4, PDGFRA, MDM2, MDM4 (Brennan et al, 2013; Mohapatra et al, 2013) are all found to be involved in tumorigenesis of a variety of cancers and are members of several signaling pathways notoriously involved in cancer.

Although these genes are highly involved in glioblastoma evolution they cannot be considered as solely astrocytoma-specific since they are malfunctioning in a great number of different cancers. EGFR (7p11.2) is one of the most renowned member of the protein kinase superfamily and a member of Ras-Raf-MEK-ERK pathway, but can also activate PI3 kinase-AKT-mTOR signaling. The gene is amplified in 40% of glioblastomas and was associated with the so called classical subtype. Nevertheless, EGFR amplification and mutations have been shown to be responsible for many other cancer types. Mohaparta et al (2011) was applying aCGH technique to biphasic anaplastic oligoastrocytoma in order to identify variations unique to each component and found that astrocytic component had many more CNAs compared to the oligodendroglial components. Although gain of chromosome 7 was present in both components amplification of EGFR and MET genes (7q31.2; PI3kinase-AKT-Mtor and Ras-Raf-MEK-ERK pathways) were confined to the astrocytic one.

Another most common amplification is of the chromosome 12 on which genes CDK4 and MDM2 reside. CDK4 (cyclin dependent kinase 4;12q14.1), yet another candidate gene for glioblastoma, is responsible for cell cycle’s G1 to S transition but is also involved in the variety of cancers. MDM2 (12q15) is an E3 ubiquitin ligase localized in the nucleus, that mediates ubiquitination of p53, leading to its degradation by the proteasome and inhibits p53- and p73-mediated cell cycle arrest and apoptosis. Similar involvement in glioblastoma displays gene MDM4 (1q32.1) (Crespo et al, 2015).

The region 10q23.31 where tumor suppressor PTEN resides is also known to be the most frequently lost in glioblastoma, but also frequently mutated or lost in a large number of other human tumors (prostate cancer, glioblastoma, endometrial, lung and breast cancer). The gene encodes a phosphatidylinositol-3,4,5-trisphosphate 3-phosphatase which contains a tensin like domain. It negatively regulates AKT/PKB signaling pathway. It is a part of the PI3K/AKT/mTOR pathway. We have observed significantly deleted region 10q24.2 distant 5814367bp from PTEN region.

Another well-known amplification event is the one on chromosome 4 (Brennan et al, 2013) where a gene for receptor tyrosine-protein kinase PDGFRA (4q12) resides. PDGFRA acts as a receptor for PDGFA, PDGFB and PDGFC growth factors necessary among other things for the growth of glial cells, too (Roskoski Jr, 2018). It has been shown that kinase PDGFRA mediates the activation of both PI3K/Akt/mTOR and Ras/Raf/MEK/ERK signaling.

Of note is our result on the deletions of loci at 9p21.3, where genes CDKN2A/CDKN2B reside, that have been identified with GISTIC as significantly deleted regions both on our total sample as well as on malignant cases only. Moreover, the region was also significant for glioblastomas. These highly recurrent homozygous deletions of CDKN2A/B genes were established (Brennan et al, 2013). CDKN2A functions as inhibitor of CDK4 kinase, which denotes this gene as tumor suppressor. Its protein can also stabilize p53 protein since it is able to sequester MDM2 ligase, a protein responsible for the degradation of p53. Adjacent to CDKN2A lays CDKN2B gene which encodes cyclin-dependent kinase inhibitor that unables the activation of CDK4 or CDK6. Both genes are involved in the G1 cell-cycle control.

Beroukhim et al (2007) report on amplifications of 4q12 and 7p11.2 (18–26% of samples) and deletions of 1p36.31 and 9p21.3 (35–49%). Their paper argues that in some cases, a high degree of amplification renders amplifications highly significant even though they occur in only 6–7% of samples. Because the background rate of deletions across the genome is higher, deletions usually must occur at higher frequencies than amplifications to attain similar levels of significance.

Roerig et al (2005) found novel sites of losses such as 15q14-q26 in anaplastic astrocytomas, which is similar to our GISTIC results encompassing total sample where region 15q15.1 was significantly deleted (even at 0.15 cutoff value). Other reported region 18q11.2–qter for secondary glioblastomas was significantly amplified in our group of pooled malignant cases (18q12.2; q=0.25), but was missing from glioblastomas probably because our collection comprised only primary glioblastomas.

Several significant aberrant regions and genes within were further investigated using functional enrichment strategies (Harris et al, 2004). According to DAVID 65 genes were assigned to a pathway or a significantly over-represented functional category. Our results on annotated genes possibly involved in astrocytoma tumors brought many candidates which we allocated to the regions identified by GISTIC. In such a manner potentially important genes in high grade samples were: SOS2, FCN3, ZNF683, FGF9, IL17D, TNFRSF19, FLT3, POLR1D, FLT1, HMGB1, genes for several interferon molecules, C1QBP, CXCL16, DHX33, GP1BA, NLRP1, P2RX1, P2RX5, CLDN7, CLEC10A, GABARAP, XAF1, DVL2, RTN4RL1, YWHAE BLNK, CHUK, ENTPD1, FGF8, HPS6, NFKB2, PIK3AP1, TAF5, TRIM8 WNT8B. Only one significantly amplified region in high grades harbored functionally relevant annotated gene - CLDN1.

Significantly deleted region suspected as an early event harbored just one functionally annotated gene - MAP1LC3A (LC3).

Heat maps revealed clear distinction between low and high grade samples showing that high grades were reflecting the general heat map images of the entire batch. Furthermore, in the majority of malignant samples systematic amplification of segments in chromosome 7 and respective deletion in chromosome 10 were evident, a pattern previously reported for glioblastoma patients (Mermel et al, 2011) as well as oral verrucous carcinoma (Samman & Sethi, 2015).

Next, we restricted our analysis to KEGG database and evidenced that only genes associated to deleted segments were significantly enriched in 18 out of 325 total Homo sapiens associated KEGG pathways. The most significantly represented pathways were PI3K- Akt, Cytokine-cytokine receptor interaction, NOD-like receptor, Jak-STAT, RIG-II-like receptor and Toll-like receptor. Also, pathways involved in viral infections and inflammation were all significantly enriched too. The enrichment map revealed roughly equal systematic involvement of all genes across the identified pathways. Probably the most intriguing of those are enrichments within HPV and Herpes simplex infection pathways as several studies indicated that the infectious agents have previously been associated with the carcinogenesis of brain and head and neck cancers (Wrensch et al, 2005; Alibek et al, 2013; Hashida et al, 2015; Strong et al, 2016; Zavala-Vega et al, 2017).

The major limitation of our study is the small number of patients in our cohort due to financial restrictions. Nevertheless, the minute CNA investigation brings important findings. We are also aware that the roles of involved genes within lost or gained regions need to be further explored by measuring their differential expression, but we must leave these experiments for future studies.

## Conclusions

As technologies progress genetic profiles and molecular findings have become recognized as potential markers of clinical distinction of tumor subtypes. Molecular characteristics are also being helpful in explaining the responses to therapy. Identifying narrow regions with altered DNA copy number is an important finding in tumor genetics, as genes mapped in these regions may represent potential candidate tumor suppressor genes and oncogenes. Despite many recent advances on the molecular biology of astrocytoma, its molecular blueprint of development and progression is still largely unexplained. Our findings contribute to better understanding of human astrocytoma genetic profile and suggest that copy number alterations play important roles in its etiology and progression. Hopefully the results of our analysis will find applicability in to clinical oncology. It would be important to validate the involvement of candidate genes employing other methods of molecular biology in further studies.

## Materials and methods

### Astrocytoma samples

The collected brain tumors were newly diagnosed and patients did not receive any treatment prior to surgical resection. The tumor samples were collected from the Departments of Neurosurgery University Hospital Centers Zagreb and “Sisters of Charity”, Zagreb, Croatia. We analyzed 14 astrocytoma samples which included two astrocytomas grade I, two astrocytomas grade II, one astrocytoma grade III and 9 glioblastomas (grade IV) (Table 9). Autologous blood samples were also obtained and analyzed for two patients (one for grade I and the other for IV). The patients had no family history of brain tumors. During the operative procedure the tumors were removed using a microneurosurgical technique after which the tissue was frozen in liquid nitrogen and transported to the laboratory, where it was immediately transferred to –80°C. The blood samples were collected in EDTA and processed immediately. Eleven patients were male, and three were female. Patient age ranged from 19 to 72 years (mean: 49.29 years; median: 50.0 years). The data on astrocytoma molecular diagnosis is shown in Table 9. Diagnosis was established on the basis of the patohistological findings by board certified neuropathologist and classified according to WHO guidelines (Louis et al, 2016). Magnetic resonance imaging (MRI) revealed the localization of astrocytic brain tumors. Ethical approvals were received from the Ethical Committees School of Medicine University of Zagreb (class number: 380-59-10106-14-55/147; Class: 641-01/14-02/01, 1.07.2014, and University Hospital Centers Zagreb (number 02/21/JG, class: 8.1.14/54-2, 23. 06. 2014.) and “Sisters of Charity” (number EP-7426/14-9,11. 06. 2014.), and the patients gave their informed consent.

**Table 9.** Clinical and epidemiological data for collected astrocytoma samples.

| Astrocytoma samples | Grades | Localization | Age at diagnosis | Sex | Molecular diagnosis |
| --- | --- | --- | --- | --- | --- |
| 1* | I | Frontoparietal R | 42 | M | ND |
| 2 | I | Occipital R | 19 | F | ND |
| 3 | II | Temporal L | 32 | M | IDH1 +; ATRX - |
| 4 | II | Insular L | 44 | M | IDH1+ |
| 5 | III | Frontal L | 24 | M | IDH1 - |
| 6* | IV | Parietal R | 68 | M | IDH1- |
| 7 | IV | Frontotemporoparietal L | 72 | M | ND |
| 8 | IV | Occipital R | 70 | M | ND |
| 9 | IV | Temporal L | 67 | M | ND |
| 10 | IV | Temporoparietal R | 55 | M | IDH1 - |
| 11 | IV | Temporoparietal L | 49 | M | IDH1 - |
| 12 | IV | Occipital L | 36 | M | IDH1 + |
| 13 | IV | Frontal R | 61 | F | IDH1 - |
| 14 | IV | Frontotemporal R | 51 | F | IDH1 - |

\*samples with analyzed blood; R=right, L= left; ND=not determined

### DNA extraction

Approximately 0.5 g of tumor tissue was homogenized with 1 ml extraction buffer (10 mM Tris HCl, pH 8.0; 0.1 M EDTA, pH 8.0; 0.5% sodium dodecyl sulfate) and incubated with proteinase K (100 μg/ml; Sigma, USA; overnight at 37°C). Phenol-chloroform extraction and ethanol precipitation followed. Blood was used to extract leukocyte DNA. Five ml of blood was lysed with 15 mL of RCLB (red blood cells lysis buffer; 155 mM NH_4_Cl; 0,1 mM EDTA; 12 mM NaHCO_3_) and centrifuged (15 min/5,000 × g) at 4°C. The pellet was further processed same as for DNA extraction from the tissue samples. Samples were purified using PCR purification kit (Qiagen). The concentrations were measured by Nanodrop and the purity of DNA was determined. Each DNA sample was analyzed on 1.5% agarose gel to assess genomic DNA intactness and the average molecular weight.

### aCGH

Array Comparative Genomic Hybridization (aCGH) was performed using SurePrint G3 Human CGH microarrays 4 × 180 K (Agilent Technologies) following the manufacturer’s instructions. Briefly, 1 μg of genomic DNAs corresponding to either a human reference control (Promega) or test samples were fragmented by heating at 95°C for 10 minutes. Fragmented DNAs were labeled with Cy3 (reference DNA) and Cy5 (test samples) fluorescent dUTP, respectively, using the SureTag Complete Labeling Kit (Agilent Technologies). Purification columns (Agilent) were used to remove the unincorporated nucleotides and dyes. The labeled samples along with human Cot-1 DNA were added together and hybridized on the array slides. Hybridizations of labeled DNAs to SurePrint G3 Human CGH Arrays (4×180K) (Agilent Technologies) were performed in a hybridization oven at 65°C at 20 rpm for 24 hours. The slide was scanned at 3 μm resolution on Agilent Microarray Scanner System (Agilent Technologies). Agilent CytoGenomics software (Agilent Technologies) was used to visualize, detect, and analyze chromosomal patterns within the microarray profiles. The true copy number variation (CNV) in the test sample was inferred from the log ratio of minimal 3 consecutive probes and gene content in the observed region. Recommended values of Agilent Technologies for log ratio in actual data are between +0.53 and -0.9.

### Bioinformatics analysis

The pipeline utilized two main computational approaches in processing the data, namely rCGH (Bioconductor package) and GISTIC (Genomic Identification of Significant Targets in Cancer) 2.0.23.

### Computation analysis of CNAs

The R package rCGH (Commo et al, 2016) was used for pre-processing, genotyping and calculation of circular binary segmentation to estimate the normalized copy number values, with circular binary segmentation carried out as implemented in the DNAcopy package (Seshan & Olshen, 2017), letting the standard deviation for segment length to be defined from the data rather than setting it to some pre-specified value. The relative log ratio centering was executed by utilizing expectation maximization algorithm thus increasing the expectation level at which a signal is being detected to its maximum. Furthermore, to increase the efficacy of the estimation process, the pipeline models the LRR distribution based on segmentation, with each segment mean and sd value (derived from probes assigned to each given segment) utilized in the process. The value was set to 0.5. Germline copy number alterations were removed from the all downstream analysis by excluding sex chromosomes.

### Functional enrichment analysis

GISTIC is software designed for discovering new cancer genes targeted by somatic copy number alterations (SCNAs) (Beroukhim et al, 2007; Como et al, 2016). To identify significantly amplified or deleted regions within a chromosome GISTIC 2.0.23 was used, by setting the confidence level to 99% for range of q-value thresholds spanning from 0.05 to 0.45 with the increment of 0.1. Focal amplification or deletion for all hg19 samples was determined by setting the broad length cutoff to 0.5, and confidence level to 0.9, with all other parameters restricted to their default values.

In order to understand the biological relevance of a list of genes obtained by GISTIC, subsequent analysis using DAVID (v6.8) (Huang et al, 2009), a functional enrichment analysis tool designed to estimate the biological relevance of a given collection of genes was performed (Kanehisa et al, 2016). The clustering algorithm in DAVID is based on the hypothesis that similar annotations should have similar gene members. It uses the Kappa statistic to measure the degree of common genes between two annotations. This is followed by heuristic clustering to group similar annotations according to Kappa values. Relevant genes were evaluated against the background consisting of only those genes queried by the microarray. For Functional Annotation Clustering considering clusters with the enrichment scores higher than 1, Fisher exact test was used to determine the significance of the obtained results utilizing two types of corrections for multiple hypothesis testing - Bonferroni (Bland & Altman, 1995) and Benjamini-Hochberg (BH) (Benjamini & Hochberg, 1995) adjusted p-values with threshold level set to α= 0.05.

Pathway enrichment analysis was performed with R packages cluster Profiler (Yu et al, 2012) and ReactomePA (Yu & He, 2016) using KEGG (Kanehisa et al, 2016, 2017) pathway database to further investigate the role of the GISTIC identified genes (associated to amplification and deletions sites) in known biological pathways considering only Benjamini-Hochberg adjusted p-values below 0.05 as significant. List of the obtained KEGG pathways together with the associated genes were mapped using R path view package (Luo and Brouwer, 2013). Bar charts were used to illustrate the number of genes that overlap in both KEGG pathways and functional annotation clusters. Moreover, dot matrices were computed to reflect the impact of genes associated with each KEGG pathway as well as enrichment maps and GSEA plots (Morgan et al, 2017).

## Acknowledgements

This work was supported by grants from Croatian Science Foundation 6625 and 9386.

## Authors’ contributions

NPŠ produced the idea, designed the study, contributed to the data collection, analysis and interpretation of the results, wrote the manuscript and revised it for important intellectual content, and approved the final version of the manuscript. AK contributed to data acquisition and analysis, performed experimental work, read the manuscript and revised it for important intellectual content. KG performed experimental work, participated in data collection, interpretation and analysis, and revision of the manuscript for important intellectual content. ML participated in date analysis and interpretation and revised the manuscript for important intellectual content; AB contributed to the data interpretation, manuscript editing, revised the manuscript for important intellectual content. RB performed biostatistical analysis and interpretation. FB contributed to data acquisition, the interpretation of the results, manuscript editing, and manuscript review. All authors read and approved the final manuscript.

## The Paper Explained

PROBLEM Which specific CNAs are attributed as early events and which are responsible for progression still remains to be fully understood. RESULTS Present study in which the grades of astrocytoma were compared with their genetic alterations brings new data to astrocytoma research this way amplifying the wide spectrum of changes. IMPACT The findings could help us identify the regions critical for the tumorigenesis, the diagnosis and the prognosis of low grade, intermediate grade and high grade astrocytic tumors.

## Expanded View information

**EV1 Fig1.** Summary of the most important identified KEGG pathways with the genes associated with deleted chromosomal regions labeled in red color.

**EV2 Fig2.** Enrichment map illustrates connectivity of significantly enriched pathways indicating the identified genes associated to deletion segments to be equally (more/less) represented in each identified pathway.

**EV3 Fig3.** Matrix layout for all genes within 18 KEGG pathways sorted by size. Dark circles in the matrix indicate functional categories with genes that are part of the intersection, that is, are associated with each pathway of the set. Bar plot above matrix depicts the number of shared genes, while the bar plot on the left shows the number of genes within a given pathway. Blue colored histograms indicate all genes are associated with deleted cytobands. S5 Table contains the description of the utilized KEGG identifiers.

**EV1 Table1.** List of genes within CNAs associated with significantly enriched functional categories as calculated using DAVID. Only significant functional enrichments are included with significance level set to Bonferroni, p<0.05. Furthermore, corresponding correction p-values were rounded up to three decimal points.

